# EmptyNN: A neural network based on positive-unlabeled learning to remove cell-free droplets and recover lost cells in single-cell RNA sequencing data

**DOI:** 10.1101/2021.01.15.426387

**Authors:** Fangfang Yan, Zhongming Zhao, Lukas M. Simon

## Abstract

Droplet-based single-cell RNA sequencing (scRNA-seq) has significantly increased the number of cells profiled per experiment and revolutionized the study of individual transcriptomes. However, to maximize the biological signal robust computational methods are needed to distinguish cell-free from cell-containing droplets. Here, we introduce a novel cell-calling algorithm called EmptyNN, which trains a neural network based on positive-unlabeled learning for improved filtering of barcodes. We leveraged cell hashing and genetic variation to provide ground-truth. EmptyNN accurately removed cell-free droplets while recovering lost cell clusters, and achieved an Area Under the Receiver Operating Characteristics (AUROC) of 94.73% and 96.30%, respectively. The comparisons to current state-of-the-art cell-calling algorithms demonstrated the superior performance of EmptyNN, as measured by the number of recovered cell-containing droplets and cell types. EmptyNN was further applied to two additional datasets and showed good performance. Therefore, EmptyNN represents a powerful tool to enhance scRNA-seq quality control analyses.

## INTRODUCTION

Droplet-based single-cell RNA sequencing (scRNA-seq) has significantly increased the number of cells profiled per experiment. As a result, droplet-based scRNA-seq enables the profiling of transcriptomes from thousands, sometimes up to several millions of cells, and provides unprecedented resolution into complex biological systems (Macosko et al., 2015; Zheng et al., 2017). In a typical experiment, the viable cells in the samples are dissociated to generate cell suspension. Every single cell in the suspension is combined with a gel bead to form a droplet containing unique barcodes. Ideally, each droplet contains one bead and one cell, which we define as a singlet (**Figure 1A**). Droplets with two or more cells are defined as doublets or multiplets. These types of droplets are cell-containing droplets. Droplets without cells are defined as empty droplets or cell-free droplets, which are expected to lack any RNA molecule.

**Figure 1.**
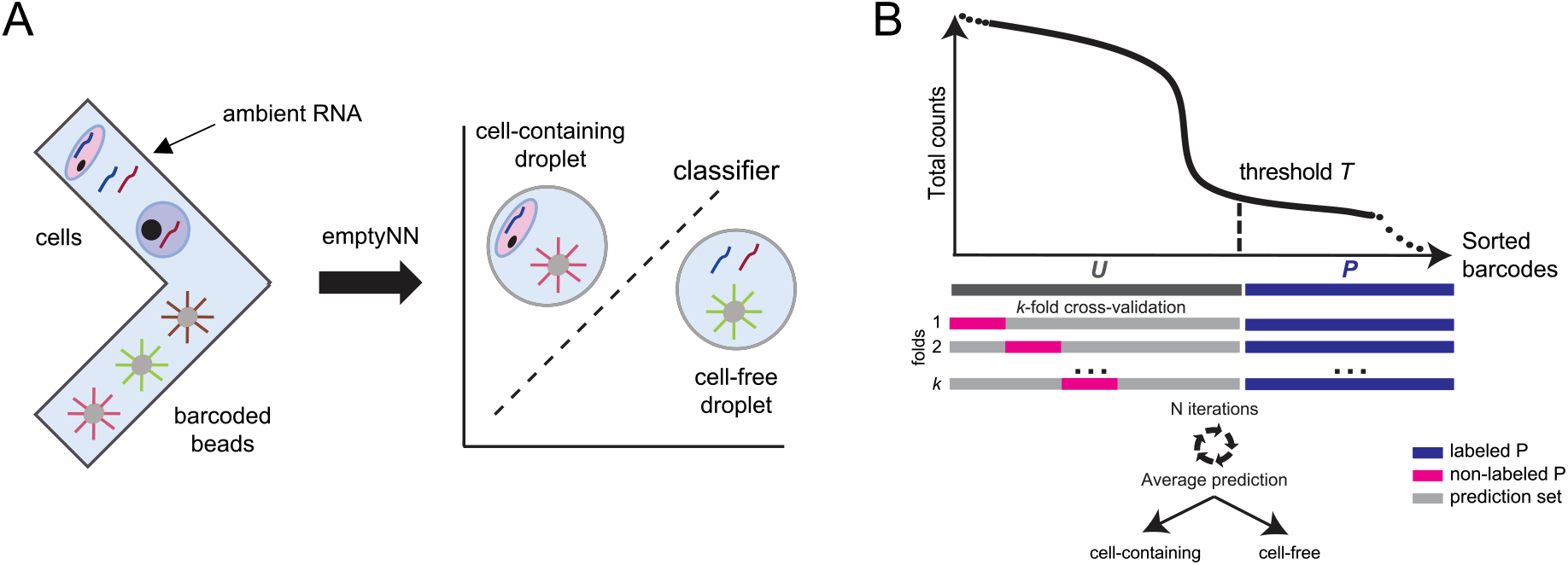
EmptyNN leverages positive unlabeled learning to classify cell-free and cell-containing droplets. (**A**) Cells and barcodes are combined in oil droplets. Some droplets may lack a cell but contain ambient RNA. The EmptyNN classifier distinguishes cell-free from cell-containing droplets. (**B**) Schematic describes the workflow of EmptyNN. The black curve represents the distribution of total counts (y-axis) across sorted barcodes (x-axis). The blue bar represents sets of barcodes with very low total counts, set *P*. The grey bar represents barcodes with higher total counts consisting of cell-containing and cell-free droplets, set *U*. EmptyNN trains a classifier, where barcodes from *P* are labeled as cell-free droplets (blue) and a fraction of barcodes from *U* is labeled as cell-containing droplets (pink). The classifier is applied to the remaining barcodes in *U* and the predictions are recorded. During each *k* fold, each barcode in *U* is predicted *k-1* times. The above process is repeated for *N* iterations (default: 10). The average prediction probability of each barcode in *U* defines each barcode as a cell-free or cell-containing droplet.

However, during cell dissociation, there exists a certain amount of subcellular debris or free-floating mRNA in the suspension, called “ambient” RNA. The ambient RNA may enter a droplet containing a barcoded bead and form a cell-free droplet (**Figure 1A**) (Young and Behjati, 2020). The ambient RNA in the cell-free droplets may be reverse transcribed into cDNA during the library preparation, which will produce unique molecular identifier (UMI) counts in the resulting gene expression matrices. Therefore, cell-free droplets are difficult to distinguish from cell-containing droplets. Failure to remove cell-free droplets may introduce spurious biological signals in the downstream analysis (Lun et al., 2019; Young and Behjati, 2020).

Computational approaches to call cells and filter barcodes in droplet-based scRNA-seq data use the following approaches. The CellRanger software from 10X Genomics (version 2 or lower) defines a cutoff based on the distribution of total counts. While very commonly used, this approach ignores any transcriptomic information. Thus, filtering barcodes solely based on the distribution of total counts may remove genuine cell-containing droplets with low RNA counts. Simultaneously, empty barcodes with high ambient RNA derived total counts may be erroneously retained.

To improve cell calling, previous work developed statistical models operating on the transcriptome profiles. Lun et al. developed EmptyDrops, which employs a Dirichlet-multinomial model to infer the transcriptome profile of cell-free droplets (Lun et al., 2019). By estimating the deviations from this profile, EmptyDrops assigns barcodes with significant deviations as cell-containing droplets. DIEM is another method that uses the multinomial mixture model along with a semi-supervised expectation-maximization algorithm to remove cell-free droplets (Alvarez et al., 2020). In addition, machine learning approaches have been successfully applied to scRNA-seq data (Angerer et al., 2017; Simon et al., 2020). One example of the application of neural networks to filter barcodes is called CellBender (Fleming et al., 2019). It uses an unsupervised deep generative model to learn the prior distribution of gene expression profiles and estimate the background RNA profile (Fleming et al., 2019).

Here, we leverage positive-unlabeled (PU) learning to train a deep neural network targeted towards cell calling. PU learning is a paradigm of semi-supervised learning, specifically designed for the case where labels of one class are available and the other class labels are uncertain (Comité et al., 1999; Denis, 1998; Letouzey et al., 2000). Contrary to standard machine learning approaches which are not suitable for such situations, PU learning takes advantage of unlabeled samples to make predictions. There are several strategies for PU learning, which involve adaptations of conventional machine learning methods, including direct application of a standard classifier (Elkan and Noto, 2008), “PU bagging” (Mordelet and Vert, 2014), and a Two-Step technique (Kaboutari et al., 2014). It was shown that PU learning achieved comparable classification performance with standard supervised machine learning approaches when applied to fully labeled data (Li and Hua, 2014).

In this manuscript, we introduce EmptyNN, a novel cell-calling algorithm that distinguishes <u>Empty</u> or cell-free droplets from cell-containing droplets by training a <u>N</u>eural <u>N</u>etwork in droplet-based scRNA-seq data. EmptyNN implements the PU learning bagging strategy and is based on the rationale that barcodes with very low total counts represent bona fide cell-free droplets. By applying EmptyNN to two ground-truth datasets and two additional datasets, we demonstrated that EmptyNN accurately discriminates cell-free and cell-containing droplets while recovering lost cell clusters with high accuracy. In our benchmarking analysis, EmptyNN outperformed current cell-calling methods CellRanger 2.0, EmptyDrops, and CellBender.

## RESULTS

In this work, we leveraged cell hashing information and genetic variation to provide ground-truth for the evaluation of our approach. We first introduced the algorithm and then compared EmptyNN to current state-of-the-art cell-calling algorithms. We conducted comprehensive benchmarking analysis. Next, we applied EmptyNN to two additional datasets and evaluated its performance. Lastly, we compared the runtime and computational requirements of the different methods.

The following section briefly describes the computational principles underlying EmptyNN. Given samples belonging to a specific class *P* and an unlabeled set *U*, which contains both *P* and *non-P* classes, the goal of PU learning is to build a binary classifier to classify *U* into two classes, *P* and *non-P* (Li and Liu, 2005; Liu et al., 2002; Mordelet and Vert, 2014). The rationale of EmptyNN is that barcodes with very low total counts represent bona fide cell-free droplets while all other barcodes could represent either cell-free or cell-containing droplets. Therefore, we defined barcodes with total counts below a user-specified threshold *T* (default: 100) as set *P* (blue) (**Figure 1B**). The remaining barcodes are defined as the unlabeled set *U* (grey), which consists of either cell-containing or cell-free droplets. Next, *U* is randomly split into *k* folds (default: 10). All barcodes in one fold are labeled as *non-P* (pink) and subsequently, a model is trained to discriminate between *P* (blue) and *non-P* (pink) barcodes. The barcodes from the remaining *k-1* folds from *U* (grey) are then predicted using the trained model and each prediction is saved. This procedure is conducted for each fold and separately repeated *N* times such that each barcode in U will be predicted *(k-1) * N* times. The average prediction is used to define each barcode as a cell-free or cell-containing droplet. We compared EmptyNN to state-of-the-art cell-calling methods CellRanger 2.0, EmptyDrops, and CellBender. More details are provided in the Methods section.

### EmptyNN removes cell-free droplets and recovers lost signal in the cell hashing dataset

To evaluate EmptyNN, we first applied it to a cell hashing dataset (Stoeckius et al., 2018). The cell hashing technology utilizes sample-specific barcodes to allow multiplexing. Cells from different donors were labeled with unique hashtag oligonucleotides (HTO) which readily separate donor samples (**Figure S1A**). Therefore, this dataset provides a unique resource to evaluate the performance of cell-calling algorithms. Specifically, the droplets that contain single or multiple HTO types were defined as singlets or doublets, respectively (**Figure S1B**). Droplets lacking a clear peak in distribution of HTO counts were defined as cell-free droplets. The barcode labels (eg. doublets, singlets, cell-free droplets) derived from the HTO information were subsequently used to evaluate the performance of EmptyNN and three competing cell-calling methods (see details in “Description of multiplexing technology and cell hashing dataset” in the Methods section).

Among the 39,842 barcodes evaluated, 20,833 barcodes (52.3%) were classified as cell-free droplets and 19,009 barcodes (47.7%) were classified as cell-containing droplets by EmptyNN. We observed that EmptyNN recovered 885 barcodes with total counts falling below the filtering threshold applied by the authors in the original study (200 in this case) (**Figure 2AB**). t-distributed stochastic neighbor embedding (t-SNE) analysis separated these recovered low total count barcodes into five unique clusters (**Figure 2C**), suggesting that they represent genuine cell-containing droplets of different cell types. Subsequent differential expression analysis revealed the cell type identities of these five clusters (**Figure 2D**). Four cell types (B, CD4, NK, and CD14 Monocytes) were present in the original study. Of note, platelets were only detected in the recovered barcodes and missed in the original study (**Figure 2C**). Since platelets contain much less RNA compared to other cell types, they are likely to be erroneously excluded from the original analysis in which filtering is only based on total counts. Indeed, the total counts of the recovered platelets were below the original filtering threshold (range: 51-196, median: 93.5).

**Figure 2.**
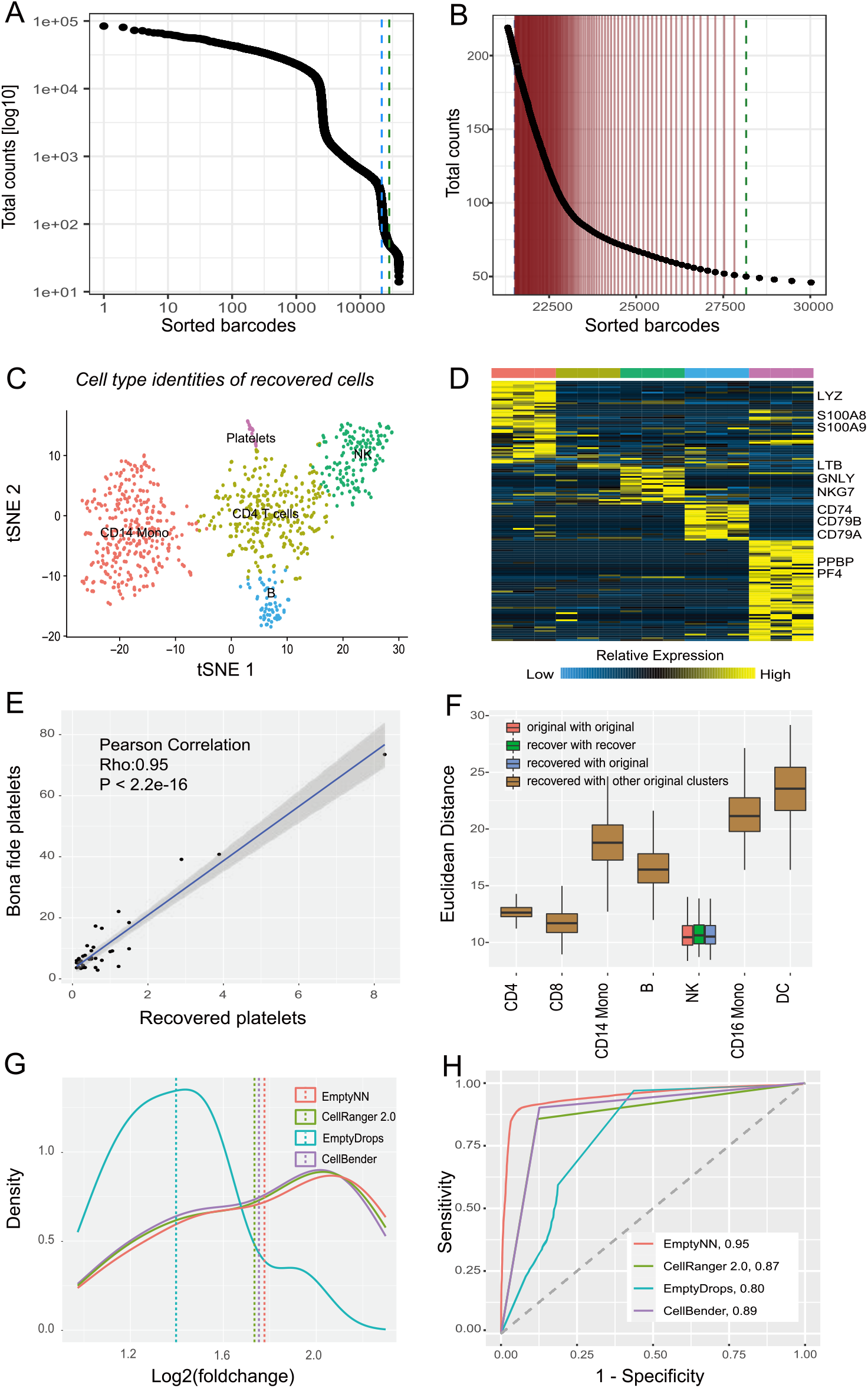
EmptyNN accurately removes cell-free droplets and recovers cell-containing droplets in the cell hashing dataset. **(A)** Barcode-rank plot shows the distribution of total UMI counts of each barcode in descending order. **(B)** A zoomed in view of barcode-rank plots highlighting barcodes with more than 50 and less than 200 total UMI counts. Vertical red lines indicate barcodes falling below the original threshold (200) but predicted to be cell-containing droplets by EmptyNN, which we referred to as “recovered cells”. (**C**) tSNE plot of the recovered cells (n = 885) shows distinct expression profiles of various cell types. (**D**) Heatmap illustrates known marker gene expression profiles derived from each recovered cluster. (**E**) Scatter plot shows significant correlation between gene expression of bona fide platelets and recovered platelets (r=0.95, p < 2.2e-16). (**F**) Boxplot shows the Euclidean distance of recovered NK cells to original clusters. (**G**) Density plot shows the differential expression signal (fold change) between “CD4 T cells” and “CD14 Monocytes” after cells called by EmptyNN, CellRanger 2.0, EmptyDrops, and CellBender, respectively. (**h**) ROC curves show the overall accuracy of different cell-calling algorithms.

To confirm the identity of the recovered platelets, we integrated an independent dataset profiling peripheral blood mononuclear cells (PBMCs) (see details in “Reference PBMC 3k dataset” in Methods section). This dataset contained a cluster of platelets which was used as a reference profile in our study. The comparison revealed a significant correlation between the gene expression profiles of bona fide platelets and our recovered platelets (Pearson correlation, Rho = 0.95, p < 2.2e-6, **Figure 2E**), demonstrating that the recovered low-count barcodes represented genuine platelets.

To further validate that the recovered barcodes represented genuine cells, we evaluated the similarity of gene expression profiles between the recovered barcodes and those present in the original study. To this end, we calculated the Euclidean distance between the following three comparisons: (a) pairwise comparison between the original cells; (b) pairwise comparison between the original and recovered cells; (c) pairwise comparison between the recovered cells. Our comparison revealed no significant differences in the distances derived from the pairwise comparison between original (red) NK cells, recovered (green) NK cells, original and recovered (blue) NK cells (One-way ANOVA, p = 0.93) (**Figure 2F**). At the same time, the recovered NK cells differed significantly from the original cells of different cell type identity (brown) (One-sided Wilcoxon rank sum test, p = 1.4e-5). These results demonstrated that the gene expression profiles of recovered NK cells did not differ from NK cells used in the original study. Analogous analyses were conducted for the other three cell types, namely B cells, CD4 T cells, and CD14 monocytes (**Figure S2**).

To benchmark our method, four cell-calling algorithms were compared: EmptyNN, CellRanger 2.0, EmptyDrops, and CellBender. By following identical preprocessing and using the identical set of highly variable genes, we performed unsupervised clustering and cell type annotation. Next, we performed differential gene expression analysis and assessed the derived expression fold changes. The rationale was that larger fold changes of well-known marker genes indicated increased biological signal. For example, when comparing CD4 T cells with CD14 monocytes, the average fold change of significant marker genes was 1.78 (std: 0.40, range: 1.04-2.30) for EmptyNN, which was higher than EmptyDrops (average: 1.40, std: 0.26, range: 0.97-1.93) (**Figure 2G**). The CellRanger 2.0 and CellBender showed similar performance to EmptyNN, with an average fold change of 1.75 (std: 0.39, range: 1.04-2.28) and 1.73 (std: 0.39, range: 1.02-2.24), respectively. These results suggested that EmptyNN slightly outperformed the other methods with respect to increasing the differential expression signal.

Furthermore, to quantitatively compare the cell-calling methods we integrated information derived from the HTO counts. As described above, the HTO counts provide singlet, doublet, and negative labels for each barcode. Barcodes labeled as singlets and doublets were classified as cell-containing barcodes and the accuracy of the four cell-calling methods was evaluated. EmptyNN achieved an AUROC (Area Under the Receiver Operating Characteristics) of 94.73% (**Figure 2H**). In contrast, CellRanger 2.0, EmptyDrops and CellBender achieved an AUROC of only 86.88%, 79.87%, and 88.80%, respectively. To visualize these results, we created a count matrix composed of all cells detected by any of these four methods. The standard analysis pipeline was applied to this count matrix, followed by the unsupervised clustering and tSNE visualization. The cell type identity of each cluster was inferred based on the HTO information and differential expression analysis results (**Figure S3AB**). Compared to CellRanger 2.0, EmptyNN retained more CD14 monocytes while discarding the ambient RNA cluster (**Figure S3CD**). Furthermore, EmptyNN retained more B cells and CD4 cells compared to EmptyDrops (**Figure S3CE**). CellBender kept most barcodes including doublet and ambient RNA clusters (**Figure S3F**). In summary, EmptyNN demonstrated superior accuracy compared to the other three cell-calling methods.

### EmptyNN accurately classifies singlets and ambiguous droplets in the multiplexed PBMC dataset

We next assessed the performance of EmptyNN in a second independent scRNA-seq dataset from Kang et al (Kang et al., 2018). Peripheral blood mononuclear cells (PBMCs) from eight individuals were pooled and then sequenced simultaneously. In the original study, the authors developed a computational tool called demuxlet, which utilizes the natural genetic variations contained in the sequencing reads to deconvolute the donor of origin for each barcode. For each barcode, demuxlet calculates the likelihood that the sequence reads originate from one or multiple individuals. Barcodes with non-discriminant probabilities are classified as ambiguous droplets which are the results of ambient RNAs from cell-free droplets. Based on this rationale, we applied demuxlet to infer the labels of each barcode and evaluated the performance of the cell-calling methods.

For a random classifier with low capacity of distinguishing between classes, the barcodes retained or discarded will have indistinguishable distributions. In contrast, for a classifier with high capacity, the distribution of retained or discarded barcodes should set apart from each other. **Figure 3A** illustrates the probability distribution of retained and discarded barcodes by EmptyNN. As expected, the median singlet probability derived from demuxlet was 1.00 (std: 0.018, range: 0.32-1.00) for retained barcodes and 0.17 (std: 0.21, range: 0.12-1.00) for discarded barcodes, suggesting its capacity of differentiating singlets and ambiguous droplets. According to the labels derived from genetic information, EmptyNN retained a total of 6,025 cell-containing droplets and achieved an AUROC of 96.30% (**Figure 3B**). The total number of retained droplets for CellRanger 2.0, EmptyDrops and CellBender was 6,061, 6,544, and 1,686, respectively. The corresponding AUROC was 82.94%, 86.43%, and 58.02%, respectively.

**Figure 3.**
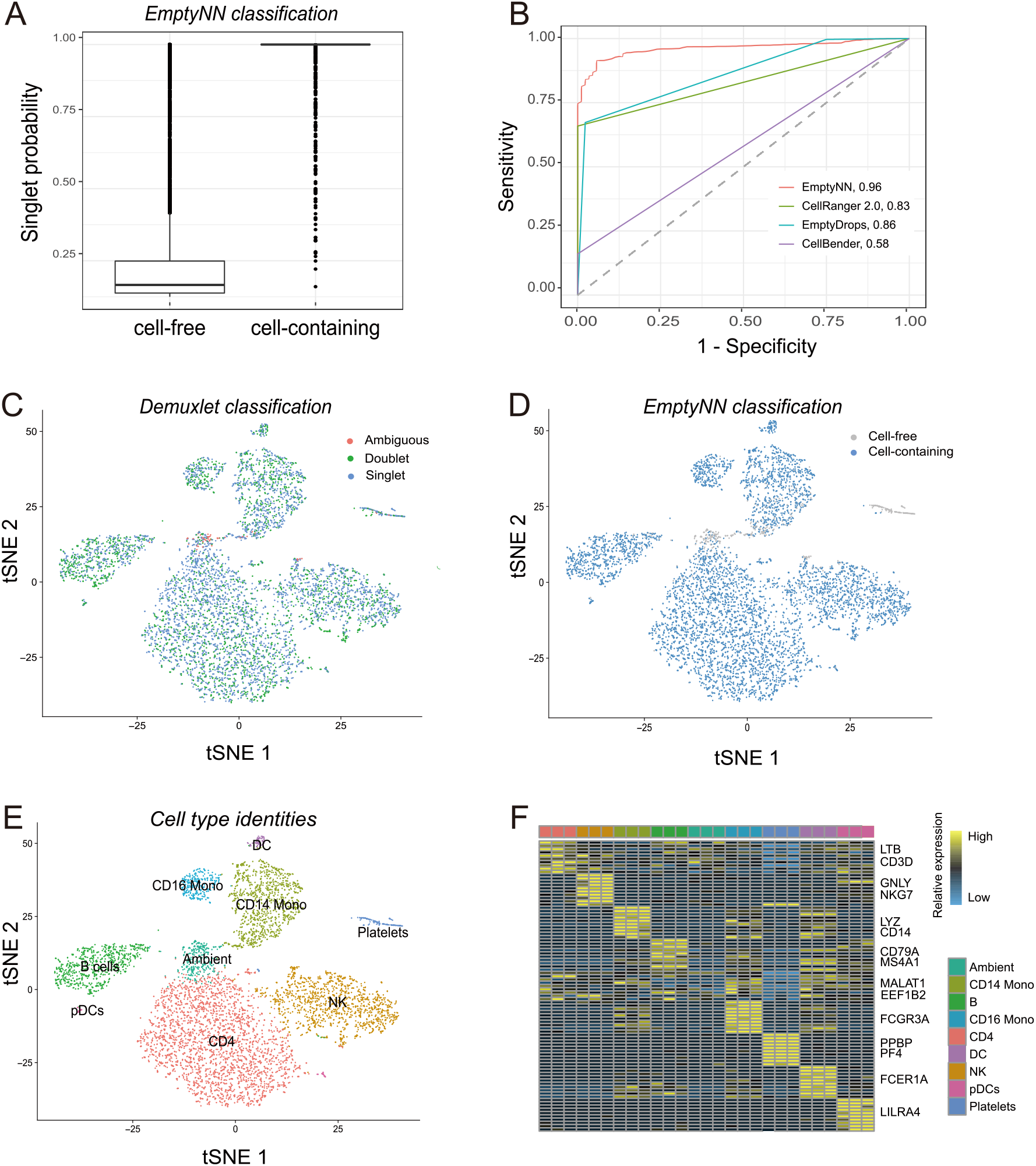
EmptyNN accurately classifies singlets and ambiguous droplets in the multiplexed PBMC dataset. **(A)** Boxplot shows the probability of being singlets for retained and discarded barcodes by EmptyNN. (**B**) ROC curve shows the performance of different algorithms. t-SNE plots visualize embedding of cells called by any of four algorithms. Points represent barcodes and are colored by **(C)** Demuxlet-derived information; **(D)** whether detected by EmptyNN; and (**E**) putative cell types. (**F**) Heatmap shows the gene expression profile for each putative cell type.

Next, we investigated the tSNE embeddings constructed from the count matrix composed of all cells detected by any of these four methods. Each droplet was labeled based on the demuxlet-derived information (**Figure 3C**). EmptyNN correctly discarded the “ambiguous” cluster (**Figure 3D**) while conserving the transcriptional profiles of the cell type identities present in the study (**Figure 3EF**). **Figure S4** shows the tSNE plots for the other methods. In summary, EmptyNN outperformed competing methods based on the accuracy inferred from genetic variation.

### EmptyNN recovers biological signals in two additional datasets

To further demonstrate the utility of EmptyNN, we analyzed two additional scRNA-seq datasets: 1) PBMC 8k dataset and 2) Neuron 900 dataset. The datasets were processed by CellRanger 2.0 and EmptyNN independently. We applied identical processing pipelines, including filtering of low-quality cells by mitochondrial fraction, normalization, highly variable gene detection, PCA analysis, clustering, and tSNE visualization (**Figure S5**). We observed that EmptyNN classified more cell-containing droplets compared to CellRanger 2.0. A critical fraction of the cell-containing droplets fell below the CellRanger 2.0 threshold and we investigated these droplets in more detail. These cell-containing droplets formed unique clusters corresponding to different cell types. In the PBMC 8k dataset, EmptyNN uniquely retained CD4 T cells, CD14 Monocytes, and platelets (**Figure 4A**), characterized by canonical marker genes, such as *LTB, CD3D, LYZ, PPBP, PF4* (**Figure 4A**). In the Neuron 900 dataset, EmptyNN recovered GABAergic, Glutamatergic, and the Non-Neuronal clusters (**Figure 4CD**). These results suggested that EmptyNN recovered genuine cell-containing droplets that otherwise would have been lost in a CellRanger 2.0 based analysis.

**Figure 4.**
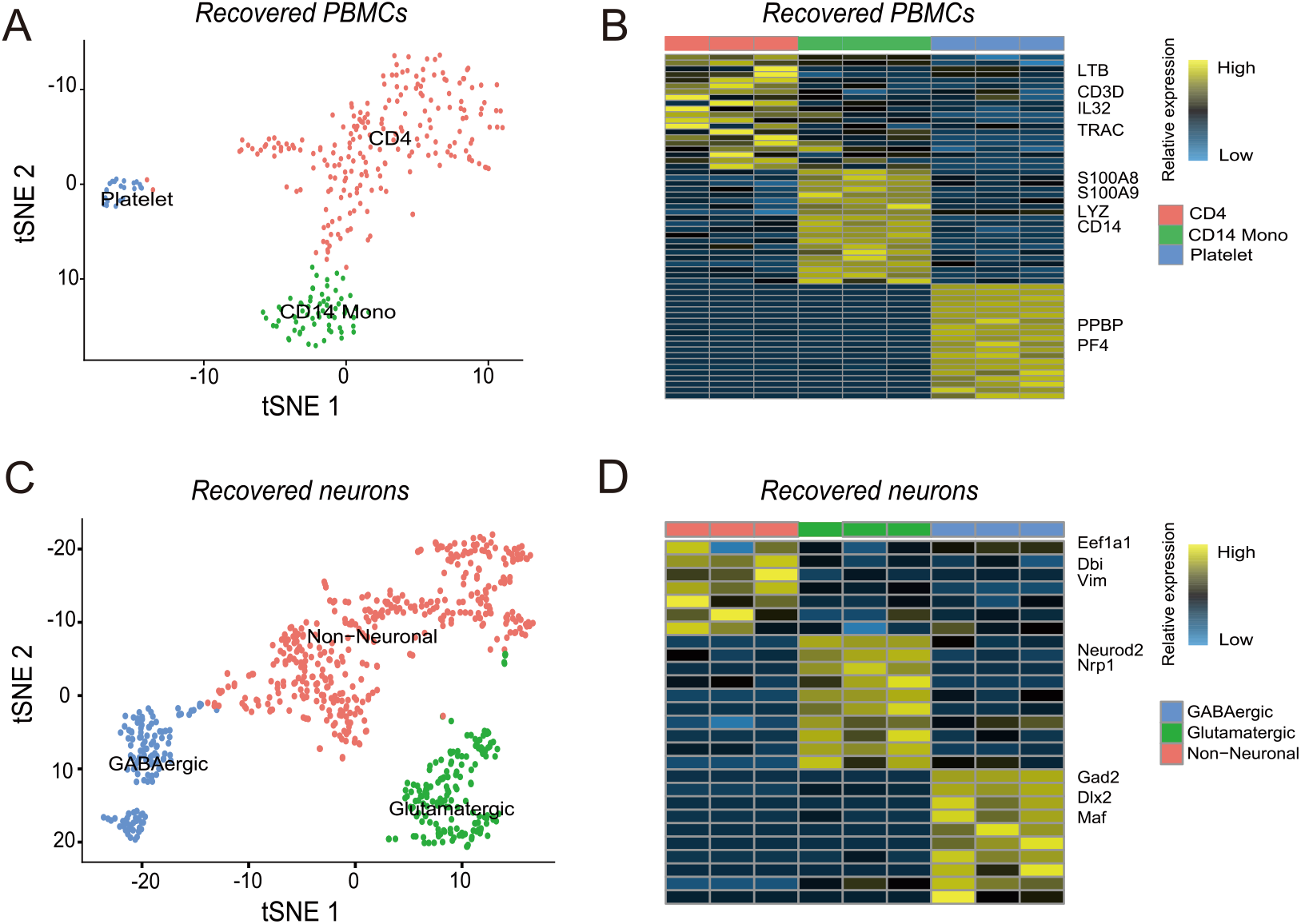
EmptyNN recovers biological signals in two additional datasets. **(A)** t-SNE plots visualize embedding of recovered barcodes by EmptyNN in the PBMC 8k dataset. Points represent barcodes and are colored by putative cell types. (**B**) Heatmap shows the gene expression profile for each cell type. **(C** and **D)** Analogous analysis and visualizations for the Neuron 900 dataset.

### Runtime comparison

Finally, we compared the cell-calling methods with respect to their runtime across all analyzed datasets (**Figure S6**). The fastest method was EmptyDrops, which finished within minutes. EmptyNN took approximately half an hour to complete. The runtime of CellBender ranged from 30 minutes (multiplexed PBMC dataset) to 17 hours (Cell hashing dataset). The results indicated that most of the methods including EmptyNN can complete the analysis within a reasonable time.

EmptyNN and EmptyDrops were implemented in the R environment and run on a standard personal computer. Based on the documentation, CellBender can be run in a CPU or GPU server. It takes approximately 30 minutes to process the full untrimmed example dataset using a CUDA-enabled GPU. In our experiments, CellBender was run on a server equipped with 24 Intel(R) Xeon(R) CPU E5-2630 v2 at 2.60GHz.

## DISCUSSION

Droplet-based scRNA-seq platforms represent a significant advancement for single-cell technologies and thus have fueled remarkable progress in our understanding of cellular systems. However, to maximize the biological signal robust computational methods are needed to distinguish cell-free from cell-containing droplets. Here, we described EmptyNN, a novel cell-calling algorithm that is based on PU learning for improved filtering of barcodes in droplet-based scRNA-seq data. We applied EmptyNN to a total of four scRNA-seq datasets and evaluated its performance.

In the cell hashing dataset, we utilized cell hashing information to assign labels (cell-free or cell-containing droplet) providing ground-truth. EmptyNN accurately classified cell-free and cell-containing droplets. We noted that EmptyNN recovered a number of barcodes with total counts falling below the filtering threshold applied by the authors in the original study. We performed independent t-SNE and differential expression analysis to infer the cell type identities of these recovered barcodes. To confirm that these barcodes represent cells, we conducted correlation analysis and Euclidean distance comparisons to demonstrate the high levels of similarity between the recovered barcodes and those present in the original study. In our benchmarking analysis, we assessed the AUROC of each cell-calling algorithm. EmptyNN achieved an AUROC of 94.73% and outperformed current state-of-the-art cell-calling algorithms. We noticed that EmptyDrops erroneously removed most CD4 and B cells. One possible explanation is that the “ambient” RNA pool is a mixture of all cell types, where the most frequent cell populations likely dominate the ambient RNA profile. EmptyDrops estimates the ambient RNA profile and assesses the deviations from this profile. Thus, the RNA profile of the most frequent cell populations may not differ sufficiently and erroneously be removed.

In the multiplexed PBMC dataset, the natural genetic variation was utilized to infer the sample identity of each barcode. The generated singlet probability of each barcode is considered as the ground-truth in this analysis. EmptyNN accurately differentiated singlets and ambiguous droplets and achieved an AUROC of 96.30%. We also applied EmptyNN to the PBMC 8k dataset and Neuron 900 dataset and demonstrated its good performance.

There are several limitations in our study. First, the key assumption of our approach is that barcodes with very low total counts represent bona fide cell-free droplets. However, this assumption may not hold when cell-containing droplets with very low total counts exist. In such cases, the user can adjust the T threshold to mitigate potential bias. Secondly, parameter selection, such as the number of cross-validation folds and the T threshold, needs to be specified manually. However, we consider our algorithm robust to various choices of k and T and plan to explore hyperparameter selection approaches in future work. Third, the retained cell-containing droplets may have a high fraction of mitochondrial reads. These low-quality cells may pass the initial filtering but can be removed in downstream analysis. Finally, ambient RNAs will remain in cell-containing droplets and contaminate the gene expression estimates. EmptyNN does not estimate corrected gene expression profiles. To correct the impact of ambient RNAs on gene expression estimates additional tools such as SoupX (Young and Behjati, 2020) need to be applied.

In summary, we introduced a novel cell-calling algorithm called EmptyNN, which is a neural network based on PU learning to discriminate cell-free from cell-containing droplets in scRNA-seq datasets. Benchmarking analysis leveraged cell hashing and genetic variation providing ground-truth, which allows for the statistical and visual comparisons of different cell-calling algorithms. We demonstrated that EmptyNN outperformed current state-of-the-art methods and accurately removed cell-free droplets while recovering genuine cells across four different datasets. We expect EmptyNN to be widely applied during the preprocessing of droplet-based scRNA-seq datasets, which will improve the downstream analysis.

## METHODS

### METHOD DETAILS

#### EmptyNN

EmptyNN is a neural network based on PU-learning. It takes the raw count matrix as input, where rows represent barcodes and columns represent genes. Only barcodes with total counts greater than 10 are included in the analysis. We define the set *P*, the bona fide empty droplets, as barcodes with total UMI counts less than threshold T (default: 100). The remaining barcodes are defined as the set *U*, the unlabeled droplets. The architecture of the neural network is composed of three layers with 128, 64, 2 neurons each. The 2000 most frequently detected genes in *P* are selected as input features of the network. The network is trained in 10 epochs using the binary cross-entropy loss function with the optimizer RMSprop. During each epoch, the training data are fed into the network in the batch size of 16. Global scaling normalization is conducted to eliminate the effect of the total counts. In this way, the neural network is forced to learn the important features in the *P* set rather than the total count difference. In each training process, the *U* set is split into *k* folds with each piece labeled as negative samples. Together with the *P* set, these split sets were fed into the neural network. The network is then used to predict the remaining *k-1* folds. The process is repeated *N* times. The barcodes in the U set will receive *(k-1) * N* predictions. Those with an average score greater than 0.5 will be labeled cell-containing droplets while those less than 0.5 will be labeled cell-free droplets. The output is a list, containing a boolean vector indicating it is a cell-containing or cell-free droplet as well as the probability of each droplet in set *U*.

#### CellRanger 2.0

CellRanger 2.0 applies an arbitrary cutoff on the total UMI counts to call cells (Zheng et al., 2017). The cutoff depends on the expected number of cells, *N*. For the top *N* barcodes, the 99th percentile of the total UMI counts is then calculated, called *m*. All barcodes with total UMI counts more than *m*/10 will be considered as cells.

#### EmptyDrops

EmptyDrops utilizes the Dirichlet-multinomial model and estimates the profile of the cell-free droplet group (Lun et al., 2019). Specifically, all barcodes were divided into three groups based on total UMI counts, including (a) cell-free droplet group or background group where total counts less than a low number (default: 100), (b) test group where total counts range from 100 and knee point, and (c) cell-containing droplet group where total counts greater than a number (default: 200). The profile of the cell-free droplet group is first estimated. Then, each barcode in the test group will be tested for deviations from this profile. Barcodes with significant deviations will be called cell-containing droplets. The EmptyDrops is implemented in the DropletUtils package (version: 1.6.1). Barcodes with FDR (False Discovery Rate) less than 0.001 will be considered as cell-containing droplets.

#### CellBender

CellBender is an unsupervised deep generative model to distinguish cell-containing droplets from cell-free ones in scRNA-seq data (Fleming et al., 2019). By utilizing a neural network, CellBender simultaneously learns the prior distribution of gene expression profiles and estimates the background RNA profile. The estimated gene expression profiles are fit with a negative binomial model to calculate the probability of each droplet containing a cell. The droplets with probability exceeding 0.5 are considered cell-containing droplets.

In our study, CellBender *remove-background* was applied to the datasets. The number of epochs was set to 150 and the learning rate was set to 1e-4 as default. The expected number of cells depends on each dataset. The barcode rank plot for each dataset was examined to determine the optimal parameter *total-droplets-included*.

### QUANTIFICATION AND STATISTICAL ANALYSIS

#### Cell hashing dataset

The cell hashing dataset from Stoeckius et al.(Stoeckius et al., 2018) utilized a multiplexing technology, which uses a unique barcoding strategy, which enables different samples to be multiplexed and sequenced together. The human peripheral blood mononuclear cells (PBMCs) from eight donors (referred to as donor A to H) were separately extracted and labeled with unique hashtag oligonucleotide (HTO). The cells from different samples were subsequently pooled and sequenced through standard scRNA-seq protocols. Both the RNA transcripts and sample unique HTO levels were obtained.

The authors applied a hard threshold to the raw count matrix and only kept barcodes with more than 200 total UMI counts. A statistical model-based strategy was developed to classify each barcode. Briefly, each HTO level was fit into a negative binomial distribution separately. The 99% quantile was used as a cutoff between “enriched” and “background”. Barcodes with HTO level above the cutoff were labeled as “positive” and barcodes below the cutoff were labeled as “negative” for that HTO. Thus, barcodes that were “positive” for only one kind of HTO were singlets. Barcodes that were “positive” for more than one kind of HTO were doublets. Barcodes that were “negative” for all HTOs were cell-free droplets. Specifically, the cutoff for HTO-A to HTO-H is 52, 75, 96, 100, 101, 128, 329, and 171, respectively.

#### Inference of droplets using hashing information

In our analysis, the raw count matrix was used, which contained 50,000 barcodes, including those barcodes with less than 200 total UMI counts. We removed barcodes without corresponding HTO information, which resulted in the exclusion of 10,158 (20.3%) barcodes. For the remaining 39,842 barcodes, the same HTO classification strategy as in the original paper was employed. In summary, barcodes were classified as 3,615 (9.07%) doublets, 19,117 (47.98%) singlets, and 17,110 (42.94%) cell-free droplets.

EmptyNN was run with 5 iterations. The threshold is set to 50. We used Seurat (version: 3.2.2) to process the EmptyNN-filtered count matrix and conduct the downstream analysis (Stuart et al., 2019). Briefly, the 1000 most highly variable genes were detected after pre-processing. Principal component analysis (PCA) was conducted to reduce dimensionality. The first ten PCs were used to calculate the neighborhood graph, followed by clustering and t-distributed stochastic neighbor embedding (t-SNE) visualization. Differential gene expression was conducted between clusters. Each cluster was labeled based on the expression of cell type marker genes. The Enrichr database (https://maayanlab.cloud/Enrichr/) served as a complementary tool for annotating cell clusters (Chen et al., 2013; Kuleshov et al., 2016).

#### Reference PBMC 3k dataset

The reference PBMC 3k dataset contained 2,700 cells in total. Standard Seurat (version: 3.2.2) pre-processing workflow was applied to this dataset, including removal of low-quality cells, normalization, feature selection, and dimension reduction. The first ten PCs were used to construct the KNN graph and cluster the cells with a resolution of 0.5. Cell markers that defined clusters were found by differential expression. There were 9 cell types in total, including naive CD4 T cells, memory CD4 cells, CD14 monocytes, CD8 T cells, CD16 Monocytes, NK, DC, and platelets. The platelet cluster served as a reference profile in our study. The mean gene expression profile of the reference and recovered platelet cluster were calculated and evaluated using Pearson correlation.

#### Comparison of Euclidean distance

The Euclidean distance was calculated based on the PC coordinates (n=50). Starting from the raw count matrix of both recovered and original cells, normalization and feature selection were conducted, followed by the PCA. The distance between any two cells was calculated using the function *dist()* in the “stats” package (version: 3.6.3). For any cell type, the pairwise distances between recovered cells and original cells were calculated. The Shapiro-Wilk test was first performed to check the normality of data. Since it did not follow a normal distribution, the Wilcoxon rank-sum tests were used to compare the means between two groups. One-way ANOVA test was conducted to compare the difference between three groups. The threshold for the p-value is set to 0.05.

#### Comparison of differential expression signal

The raw count matrix was processed by each cell-calling algorithm independently. The filtered matrix was normalized and scaled using the same set of highly variable genes. We next performed unsupervised clustering and cell type annotation. The differential expression analysis was conducted using *FindMarkers()* in the Seurat package. The expression fold changes of top significant genes between two clusters were extracted and compared.

#### Comparison of overall accuracy

We utilized the “pROC” package (version: 1.16.2) in R to calculate the overall accuracy of each cell-calling algorithm. The true labels came from cell hashing information or genetic variation. EmptyNN and EmptyDrops output the probability for each predicted barcode while CellRanger 2.0 and CellBender output the boolean vector showing cell-free droplet or not.

#### Multiplexed PBMC dataset

We first downloaded the genome-aligned bam file from the Sequence Read Archive (SRR5398237). The “bamtofastq” tool (1.3.2, https://support.10xgenomics.com/docs/bamtofastq) was used to convert the bam file to FASTQ files which were subsequently used as input to the CellRanger 2.0 pipeline. The pipeline was run with default parameters to generate the unfiltered count matrix (n = 145,549 barcodes).

#### Inference of droplets using genetic variation

To assign labels to each barcode, we used demuxlet (Kang et al., 2018), a tool to deconvolute pooled sample identities based on natural genetic variation. Demuxlet requires two inputs: 1) a BAM file containing aligned reads, and 2) a VCF file containing the genotype of each pooled sample. The output contains the most likely sample identity for each barcode in form of probabilities. The merged VCF file containing all samples was downloaded from the demuxlet github repository (https://github.com/yelabucsf/demuxlet_paper_code/). Demuxlet was run via Docker using default parameters. The output contained 5,845 singlets, 2,401 doublets, and 31,700 ambiguous droplets.

#### PBMC 8k dataset

The PBMC 8k dataset contains peripheral blood mononuclear cells (PBMCs) from a healthy donor. There are a total of 409,508 barcodes with non-zero total counts in the raw count matrix. The EmptyNN was applied to the dataset with default parameters. There were 9,685 barcodes predicted to be cell-containing droplets. The Cell Ranger 2.0 filtration results in 8,381 barcodes.

#### Neuron 900 dataset

The samples for Neuron 900 dataset come from a combined cortex, hippocampus, and subventricular zone of an E18 mouse. The raw count matrix contains 737, 280 barcodes, with 231,912 (31.46%) barcodes having at least one gene with observed expression. The EmptyNN was applied to the dataset with the threshold set to 100. After 5 iterations, there were 2899 barcodes predicted to be cell-containing droplets. In the Cell Ranger 2.1.0 filtered data matrix, there were 931 barcodes. The EmptyDrops was run with default parameters. There were 1930 barcodes with FDR less than 0.001 and considered as cell-containing droplets.

## ACKNOWLEDGMENTS

This research was partially supported by the Cancer Prevention and Research Institute of Texas grant (CPRIT RP180734). Z.Z. was partially supported by the National Institutes of Health grant (R01LM012806 and R01DE029818). The funder had no role in the study design, data collection, and analysis, decision to publish, or preparation of the manuscript.

## AUTHOR CONTRIBUTIONS

Conceptualization, L.M.S.; Methodology, L.M.S. and F.Y.; Formal Analysis, F.Y. and L.M.S.; Data curation: F.Y.; Writing – Original Draft, F.Y. and L.M.S.; Writing – Review & Editing, L.M.S. and Z.Z.; Visualization: F.Y. and L.M.S.; Funding Acquisition, Z.Z.; Supervision, L.M.S. and Z.Z.

## DATA AVAILABILITY

The BAM file of the multiplexed PBMC dataset was obtained from the SRA website (https://www.ncbi.nlm.nih.gov/sra). All other datasets were obtained from the GEO database (https://www.ncbi.nlm.nih.gov/geo/). Detailed information can be found in Table S1. The source code and tutorials are freely available at https://github.com/lkmklsmn/emptynn.

## SUPPLEMENTAL INFORMATION

**Table S1.** Description of datasets used in this study.

| **Dataset** | **Description** | **Technologies** | **Web link** |
| --- | --- | --- | --- |
| Cell hashing dataset | Human PBMCs from 8 donors | CITE-seq (10x Genomics) | https://www.ncbi.nlm.nih.gov/geo/query/acc.cgi?acc=GSE108313 |
| Reference 3k PBMC dataset | Human PBMCs from one healthy individual | 10x Genomics | https://support.10xgenomics.com/single-cell-gene-expression/datasets/1.1.0/pbmc3k |
| Multiplexed PBMC dataset | Human PBMCs from 8 individuals | 10x Genomics | CellRanger 2.0 filtered count matrix:<br>https://www.ncbi.nlm.nih.gov/geo/query/acc.cgi?acc=GSM2560247<br>BAM file:<br>https://sra-pub-src-1.s3.amazonaws.com/SRR5398237/C.merged.bam.1<br>merged VCF file:<br>https://github.com/yelabucsf/demuxlet_paper_code/tree/master/fig2 |
| 10x PBMC 8k dataset | Human PBMCs from a healthy donor | 10x Genomics | https://support.10xgenomics.com/single-cell-gene-expression/datasets/2.1.0/pbmc8k |
| 10x Neuron 900 dataset | Mouse brain cells from stage E18 | 10x Genomics | https://support.10xgenomics.com/single-cell-gene-expression/datasets/2.1.0/neurons_900 |

PBMC: peripheral blood mononuclear cell;
CITE-seq: Cellular Indexing of Transcriptomes and Epitopes by Sequencing;
E18: embryonic day 18.

**Figure S1.**
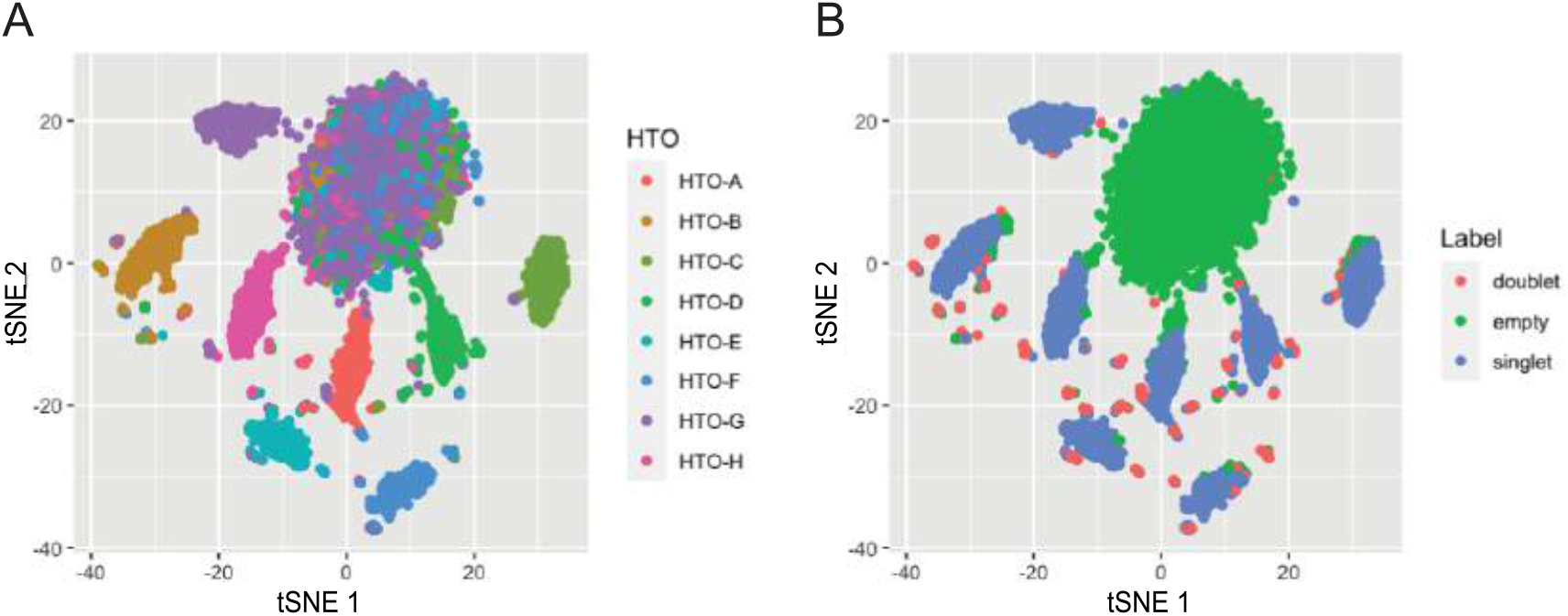
Cell hashing experiment provides ground truth definitions for cell-containing and cell-free droplets. Scatter plots illustrate t-SNE embedding of the HTO count matrix colored by (**A**) HTO samples and (**B**) barcode labels. In (B), it readily separates cell-free (green) from cell-containing (red and blue) barcodes. (**C**) t-SNE embedding of the RNA expression matrix reveals PBMC cell type identities.

**Figure S2.**
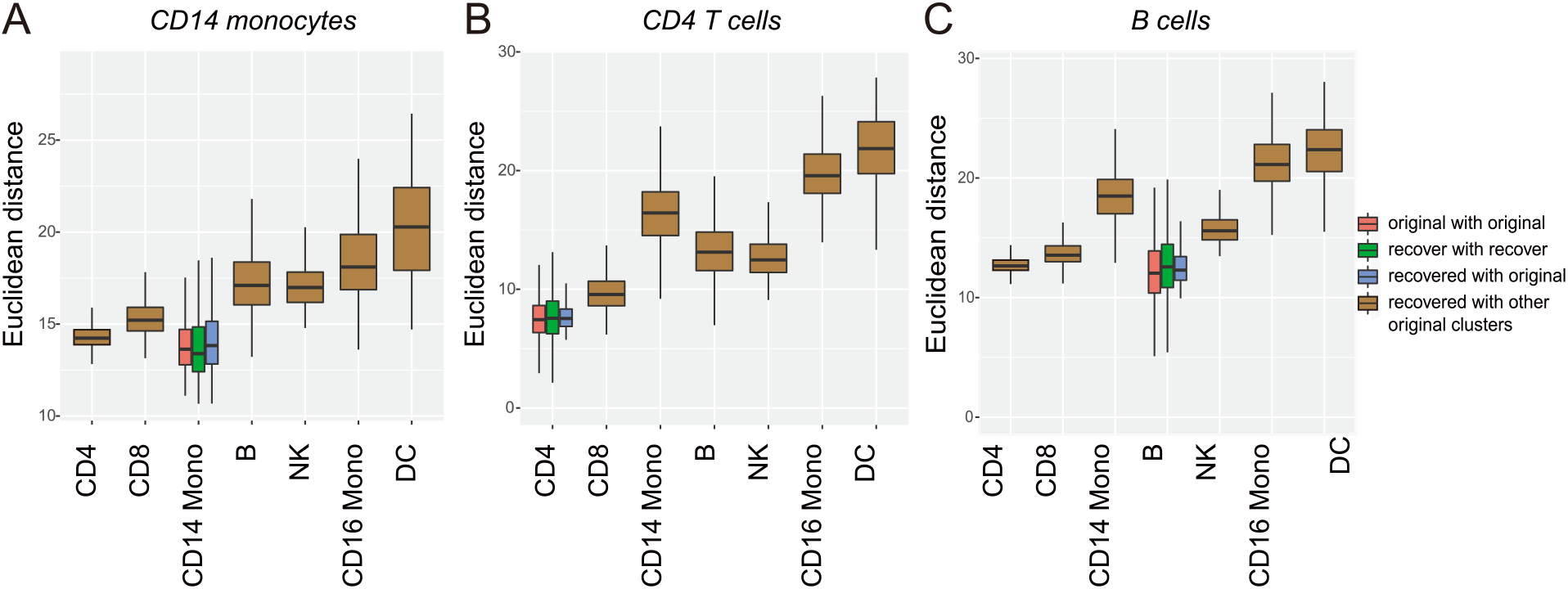
Boxplots depict the similarity between recovered barcodes and those present in original study. Boxplots show the Euclidean distance of (**A**) recovered CD14 monocytes, (**B**) recovered CD4 T cells, and (**C**) recovered B cells to original clusters.

**Figure S3.**
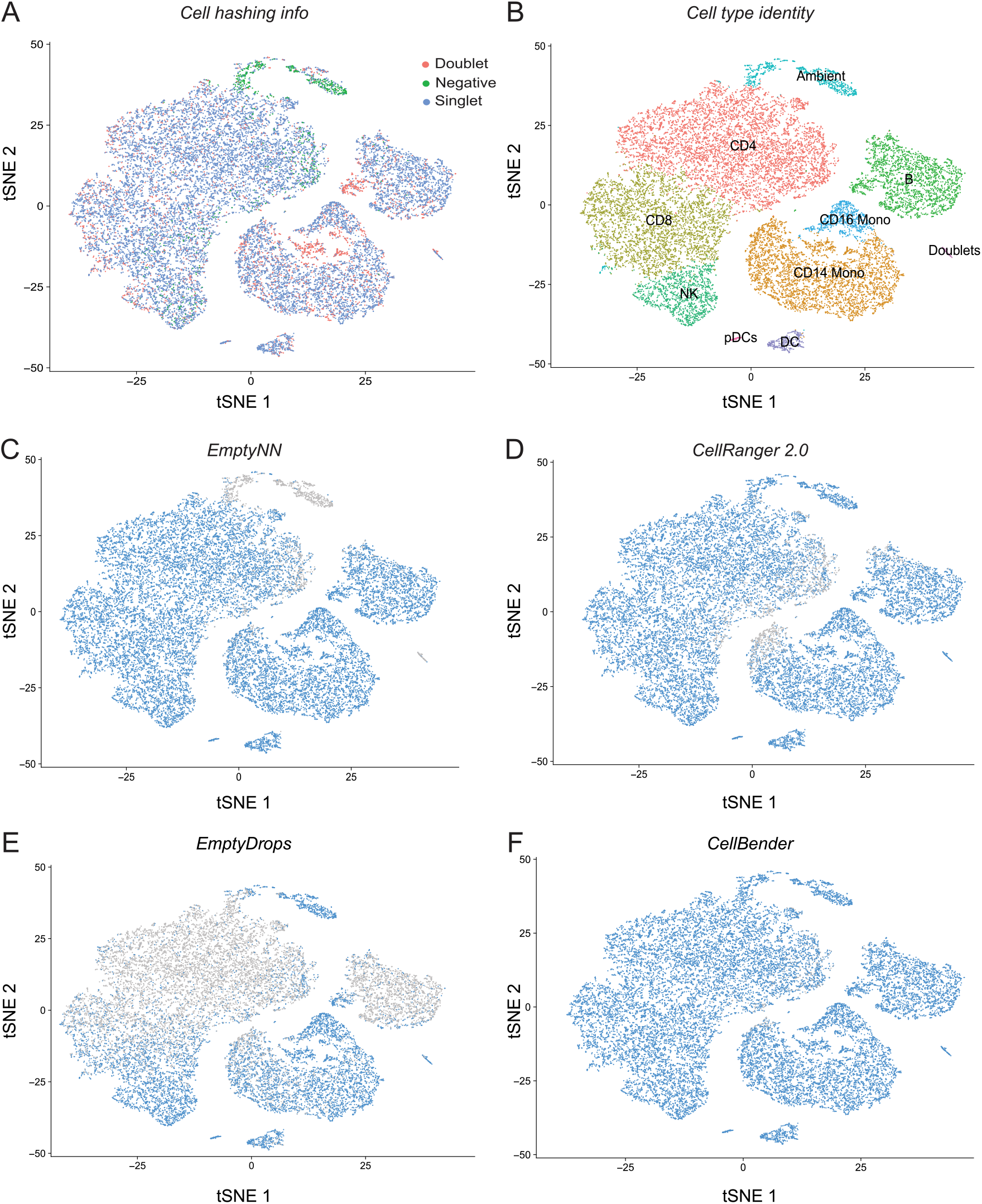
tSNE plot illustrates all cells called by any of these four cell-calling methods in the cell hashing dataset. Each dot denotes a cell and is colored based on (**A**) inferred identity (doublet, singlet, cell-free droplet) derived from hashing information; (**B**) inferred cell type derived from the transcriptional profile. The following four panels show whether the cell is called by (**C**) EmptyNN; (**D**) CellRanger 2.0; (**E**) EmptyDrops; or (**F**) CellBender.

**Figure S4.**
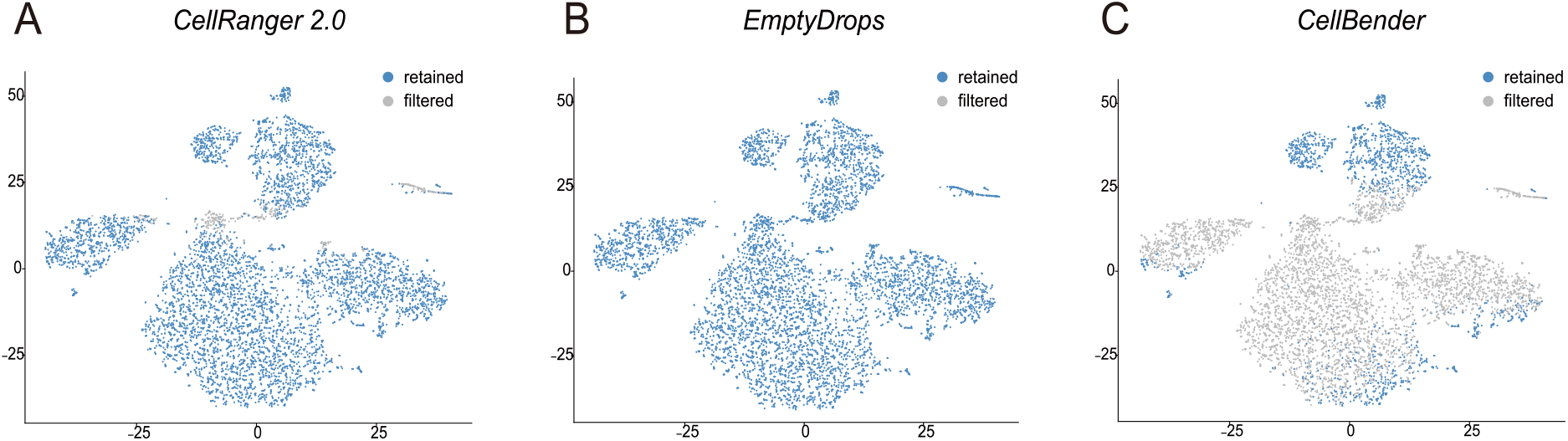
t-SNE plot highlights cells called by the other three cell-calling methods in the multiplexed PBMC dataset. Points represent barcodes colored by classifications derived from **(A)** CellRanger 2.0, (**B**) EmptyDrops, and (**C**) CellBender.

**Figure S5.**
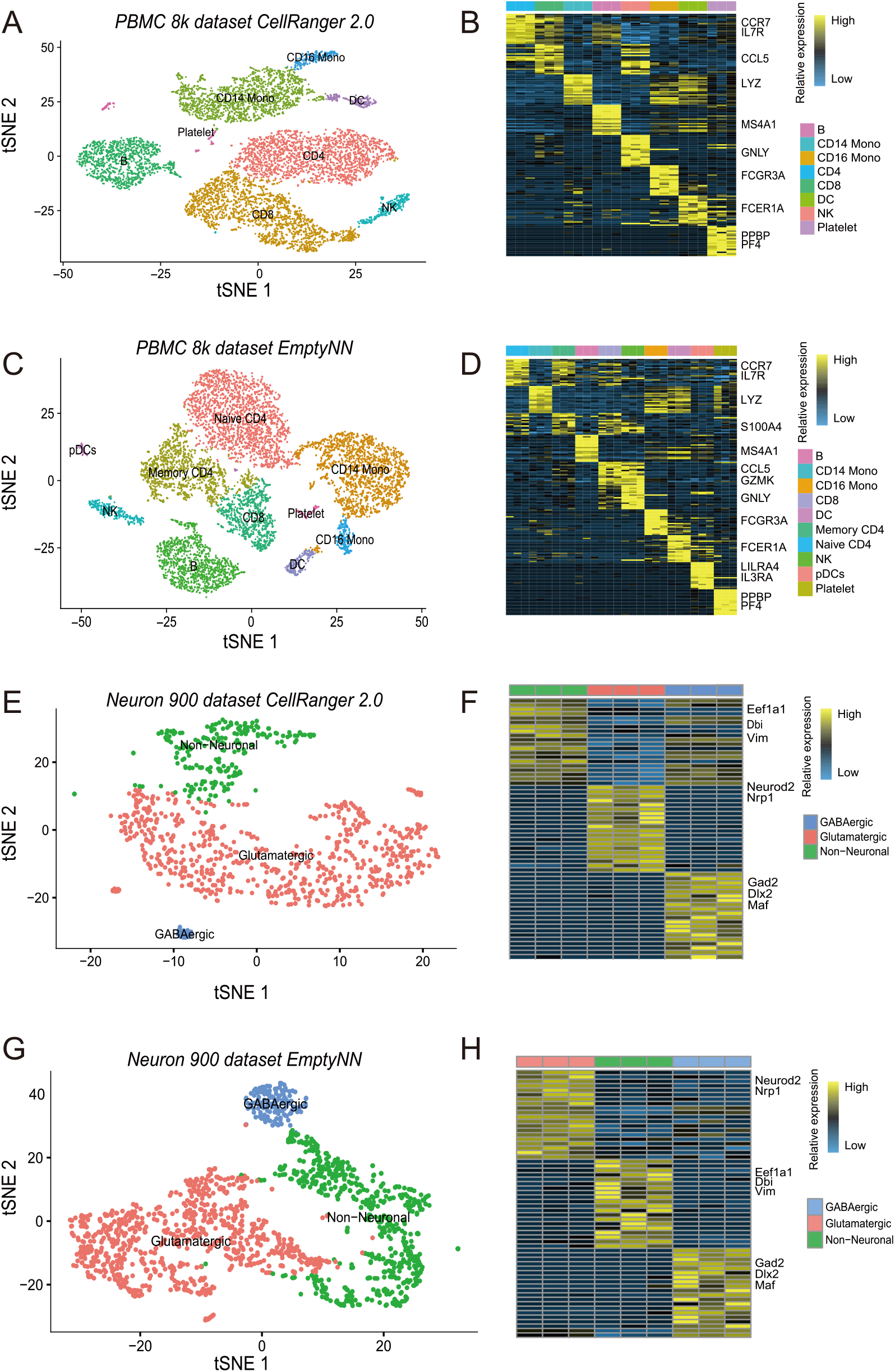
EmptyNN recovers biological signals in two additional datasets. t-SNE visualization and corresponding heatmap of the PBMC 8k dataset processed independently by CellRanger 2.0 (**A,B**) and EmptyNN (**C,D**). t-SNE visualization and corresponding heatmap of the Neuron 900 dataset processed independently by CellRanger 2.0 (**E,F**) and EmptyNN (**G,H**).

**Figure S6.**
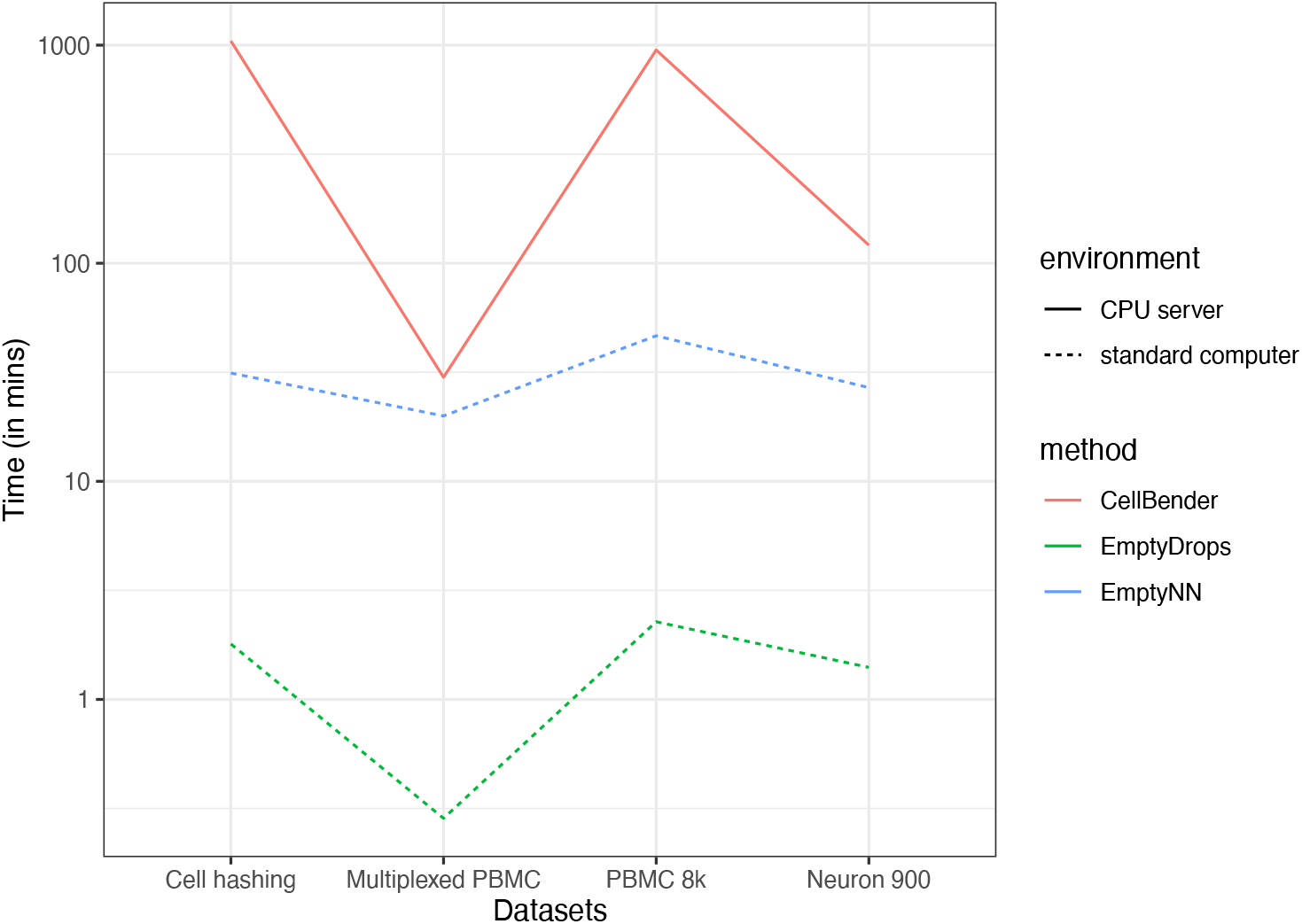
Runtime comparison. The line plot shows runtime in minutes (y-axis) across different datasets (x-axis). Color represents the different cell-calling algorithms. Line type represents the running environment.

## REFERENCES

Alvarez, M., Rahmani, E., Jew, B., Garske, K.M., Miao, Z., Benhammou, J.N., Ye, C.J., Pisegna, J.R., Pietiläinen, K.H., Halperin, E., et al. (2020). Enhancing droplet-based single-nucleus RNA-seq resolution using the semi-supervised machine learning classifier DIEM. Sci. Rep. 10, 11019.

Angerer, P., Simon, L., Tritschler, S., Alexander Wolf, F., Fischer, D., and Theis, F.J. (2017). Single cells make big data: New challenges and opportunities in transcriptomics. Current Opinion in Systems Biology 4, 85–91.

Chen, E.Y., Tan, C.M., Kou, Y., Duan, Q., Wang, Z., Meirelles, G.V., Clark, N.R., and Ma’ayan, A. (2013). Enrichr: interactive and collaborative HTML5 gene list enrichment analysis tool. BMC Bioinformatics 14, 128.

Comité, F.D., De Comité, F., Denis, F., Gilleron, R., and Letouzey, F. (1999). Positive and Unlabeled Examples Help Learning. Lecture Notes in Computer Science 219–230.

Denis, F. (1998). PAC Learning from Positive Statistical Queries. In Algorithmic Learning Theory, (Springer Berlin Heidelberg), pp. 112–126.

Elkan, C., and Noto, K. (2008). Learning classifiers from only positive and unlabeled data. In Proceedings of the 14th ACM SIGKDD International Conference on Knowledge Discovery and Data Mining, (New York, NY, USA: Association for Computing Machinery), pp. 213–220.

Fleming, S.J., Marioni, J.C., and Babadi, M. (2019). CellBender remove-background: a deep generative model for unsupervised removal of background noise from scRNA-seq datasets.

Kaboutari, A., Bagherzadeh, J., and Kheradmand, F. (2014). An evaluation of two-step techniques for positive-unlabeled learning in text classification. Int. J. Comput. Appl. Technol. Res 3, 592–594.

Kang, H.M., Subramaniam, M., Targ, S., Nguyen, M., Maliskova, L., McCarthy, E., Wan, E., Wong, S., Byrnes, L., Lanata, C.M., et al. (2018). Multiplexed droplet single-cell RNA-sequencing using natural genetic variation. Nat. Biotechnol. 36, 89–94.

Kuleshov, M.V., Jones, M.R., Rouillard, A.D., Fernandez, N.F., Duan, Q., Wang, Z., Koplev, S., Jenkins, S.L., Jagodnik, K.M., Lachmann, A., et al. (2016). Enrichr: a comprehensive gene set enrichment analysis web server 2016 update. Nucleic Acids Res. 44, W90–W97.

Letouzey, F., Denis, F., and Gilleron, R. (2000). Learning From Positive and Unlabeled Examples. In Algorithmic Learning Theory, (Springer Berlin Heidelberg), pp. 71–85.

Li, C., and Hua, X.-L. (2014). Towards Positive Unlabeled Learning for Parallel Data Mining: A Random Forest Framework. In Advanced Data Mining and Applications, (Springer International Publishing), pp. 573–587.

Li, X.-L., and Liu, B. (2005). Learning from Positive and Unlabeled Examples with Different Data Distributions. In Machine Learning: ECML 2005, (Springer Berlin Heidelberg), pp. 218–229.

Liu, B., Lee, W.S., Yu, P.S., and Li, X. (2002). Partially supervised classification of text documents. In ICML, pp. 387–394.

Lun, A.T.L., Riesenfeld, S., Andrews, T., Dao, T.P., Gomes, T., participants in the 1st Human Cell Atlas Jamboree, and Marioni, J.C. (2019). EmptyDrops: distinguishing cells from empty droplets in droplet-based single-cell RNA sequencing data. Genome Biol. 20, 63.

Macosko, E.Z., Basu, A., Satija, R., Nemesh, J., Shekhar, K., Goldman, M., Tirosh, I., Bialas, A.R., Kamitaki, N., Martersteck, E.M., et al. (2015). Highly Parallel Genome-wide Expression Profiling of Individual Cells Using Nanoliter Droplets. Cell 161, 1202–1214.

Mordelet, F., and Vert, J.-P. (2014). A bagging SVM to learn from positive and unlabeled examples. Pattern Recognit. Lett. 37, 201–209.

Simon, L.M., Yan, F., and Zhao, Z. (2020). DrivAER: Identification of driving transcriptional programs in single-cell RNA sequencing data. Gigascience 9.

Stoeckius, M., Zheng, S., Houck-Loomis, B., Hao, S., Yeung, B.Z., Mauck, W.M., Smibert, P., and Satija, R. (2018). Cell Hashing with barcoded antibodies enables multiplexing and doublet detection for single cell genomics. Genome Biology 19.

Stuart, T., Butler, A., Hoffman, P., Hafemeister, C., Papalexi, E., Mauck, W.M., 3rd, Hao, Y., Stoeckius, M., Smibert, P., and Satija, R. (2019). Comprehensive Integration of Single-Cell Data. Cell 177, 1888–1902.e21.

Young, M.D., and Behjati, S. (2020). SoupX removes ambient RNA contamination from droplet based single-cell RNA sequencing data. GigaScience 9, giaa151.

Zheng, G.X.Y., Terry, J.M., Belgrader, P., Ryvkin, P., Bent, Z.W., Wilson, R., Ziraldo, S.B., Wheeler, T.D., McDermott, G.P., Zhu, J., et al. (2017). Massively parallel digital transcriptional profiling of single cells. Nat. Commun. 8, 14049.

